# High resolution dynamic mapping of the *C. elegans* intestinal brush border

**DOI:** 10.1101/2021.06.14.448317

**Authors:** Aurélien Bidaud-Meynard, Flora Demouchy, Ophélie Nicolle, Anne Pacquelet, Shashi Kumar Suman, Camille Plancke, François Robin, Grégoire Michaux

**Author notes:** These authors contributed equally to this work (Flora Demouchy, Ophélie Nicolle, Anne Pacquelet).

## Abstract

The intestinal brush border is made of an array of microvilli that increases the membrane surface area for nutrient processing, absorption, and host defence. Studies on mammalian cultured epithelial cells uncovered some of the molecular players and physical constrains required to establish this apical specialized membrane. However, the building and maintenance of a brush border *in vivo* has not been investigated in detail yet. Here, we combined super-resolution imaging, transmission electron microscopy and genome editing in the developing nematode C. *elegans* to build a high-resolution and dynamic localization map of known and new markers of the brush border. Notably, we show that microvilli components are dynamically enriched at the apical membrane during microvilli outgrowth and maturation but become highly stable once microvilli are built. This new mapping tool will be instrumental to understand the molecular processes of microvilli growth and maintenance *in vivo* as well as the effect of genetic perturbations, notably in the context of disorders affecting brush border integrity.

## Introduction

The intestinal exchange surface required for efficient nutrient absorption is reached through three consecutive morphogenic processes in mammals: elongation and folding of the intestinal tube, generation of villus protrusions and establishment of microvilli at the surface of each enterocyte, which together amplify the tube area nearly 100-times (Helander and Fandriks, 2014). Generation of microvilli in mammals occurs during enterocyte differentiation along the crypt-villus axis with the nucleation of actin filaments anchored on an F-actin- and intermediate filament-based terminal web. These core-actin bundles are then organized into a well-ordered and tightly packed array by various actin crosslinking and bundling factors, such as villin, espin and plastinl/fimbrin (Sauvanet et al., 2015). Recent studies in epithelial cell lines identified new functional players, such as IRTKS or myosin la/6/7b (Crawley et al., 2014a, Postema et al., 2018), and new mechanisms of brush border assembly and maintenance by vesicular trafficking (Vogel et al., 2015), microvilli motility, contraction and clustering (Meenderink et al., 2019, Chinowsky et al., 2020) or intermicrovillar protocadherin bridges (Crawley et al., 2014b). Additionally, recent use of live imaging revealed some of the key initiation and maturation steps of microvilli biogenesis in cell lines (Gaeta et al., 2021). However, a description of brush border formation and maintenance *in vivo* is lacking.

The soil nematode C. *elegans* has been widely used as a *in vivo* model of intestinal luminogenesis, polarity, and host defence (Zhang et al., 2013, Zhang and Hou, 2013, Sato et al., 2014). Intestinal organogenesis in C. *elegans* encompasses cell division and intercalation steps from the E blastomere ancestor to form a primordium containing two rows of eight cells (E16 stage) which ends up, after a last round of division, with twenty cells arranged into nine rings (or *ints)* forming a ellipse-shaped tube that runs along the whole body of the worm (Leung et al., 1999, Asan et al., 2016). Contrary to the 3-5-days living mammalian enterocytes that arise from the proliferation of crypt based columnar stem cells, the C. *elegans* intestine is composed of perennial epithelial cells (Walton et al., 2016, McGhee, 2007). Their polarization begins at the E16 stage and encompasses cellular components relocalization and cell shape changes (Leung et al., 1999) as well as the apical accumulation of the polarity determinant PAR-3 which recruits the other members of the PAR polarity complex (Feldman and Priess, 2012, Achilleos et al., 2010). Luminogenesis concomitantly occurs at the E16 stage with the formation of apical cavities at the midline that ultimately form a lumen, a process that probably involves vesicular trafficking (Leung et al., 1999). C. *elegans* enterocytes displays a brush border that is structurally similar to that ofmammals (Leung et al., 1999, Geisler et al., 2019, Bidaud-Meynard et al., 2019) and relies on some of the same structural components. Indeed, C. *elegans* microvilli are made of F-actin core bundles, notably the intestinal-specific isoform of actin *act-5,* whose depletion induces a circular lumen with sparse and defective microvilli (MacQueen et al., 2005). Several F-actin regulators have been shown to be essential for C. *elegans* brush border integrity, such as *erm-1,* the only ortholog of the Ezrin/Radixin/Moesin family (Gobel et al., 2004, Van Furden et al., 2004) and the actin capping factor EPS-8 (Croce et al., 2004). As in mammals, these microvilli are anchored on a terminal web made ofa network of F-actin and various intermediate filament isoforms forming an electron-dense belt named as endotube, in which IFB-2 plays a major role (Geisler et al., 2020, Bossinger et al., 2004). Hence, the structural and biochemical similarity with mammals make C. *elegans* an appropriate model to study the biogenesis of microvilli *in vivo.*

Most of the studies in C. *elegans* focused on early polarization steps and the formation of the brush border was only studied by Transmission Electron Microscopy (TEM), which provides ultrastructural data but lacks the dynamics and whole organ context. Here, we combined super-resolution and quantitative live microscopy, TEM and fluorescence recovery after photobleaching data to characterize the recruitment and dynamics of endogenously tagged markers during the establishment of the brush border *in vivo* in C. *elegans.*

## Results and discussion

### TEM analysis of the brush border establishment in C. elegans developing embryo

To first characterize the development of the brush border *in vivo,* embryos and larvae at various developmental stages were analysed by TEM (Nicolle et al., 2015). We observed that the intestinal lumen starts to open at the comma stage and progressively expands to reach the renown elliptic shape in larvae (Fig. 1A-B). At the apex, the first microvilli-like membrane extensions were observed at the 1.5-fold stage and started to cover the apical membrane, with a disorganized pattern, at the 2.5-fold stage, to finally form a regular brush border from the 3-fold stage (Fig. 1A). Measurements suggest a relatively continuous increase in microvilli density, length and width (Figs 1A-E and S1A-C) which implies a gradual maturation of the brush border as well as *de nova* growth of microvilli throughout development to fill in the membrane added during intestinal surface expansion (Fig. 1A, arrows). Finally, transversal imaging of the brush border allowed to measure the distance between microvilli edges and centres (76,0 ± 1,1 nm and 203,2 ± 2,0 nm, respectively) (Fig. 1F-G).

**Figure 1.**
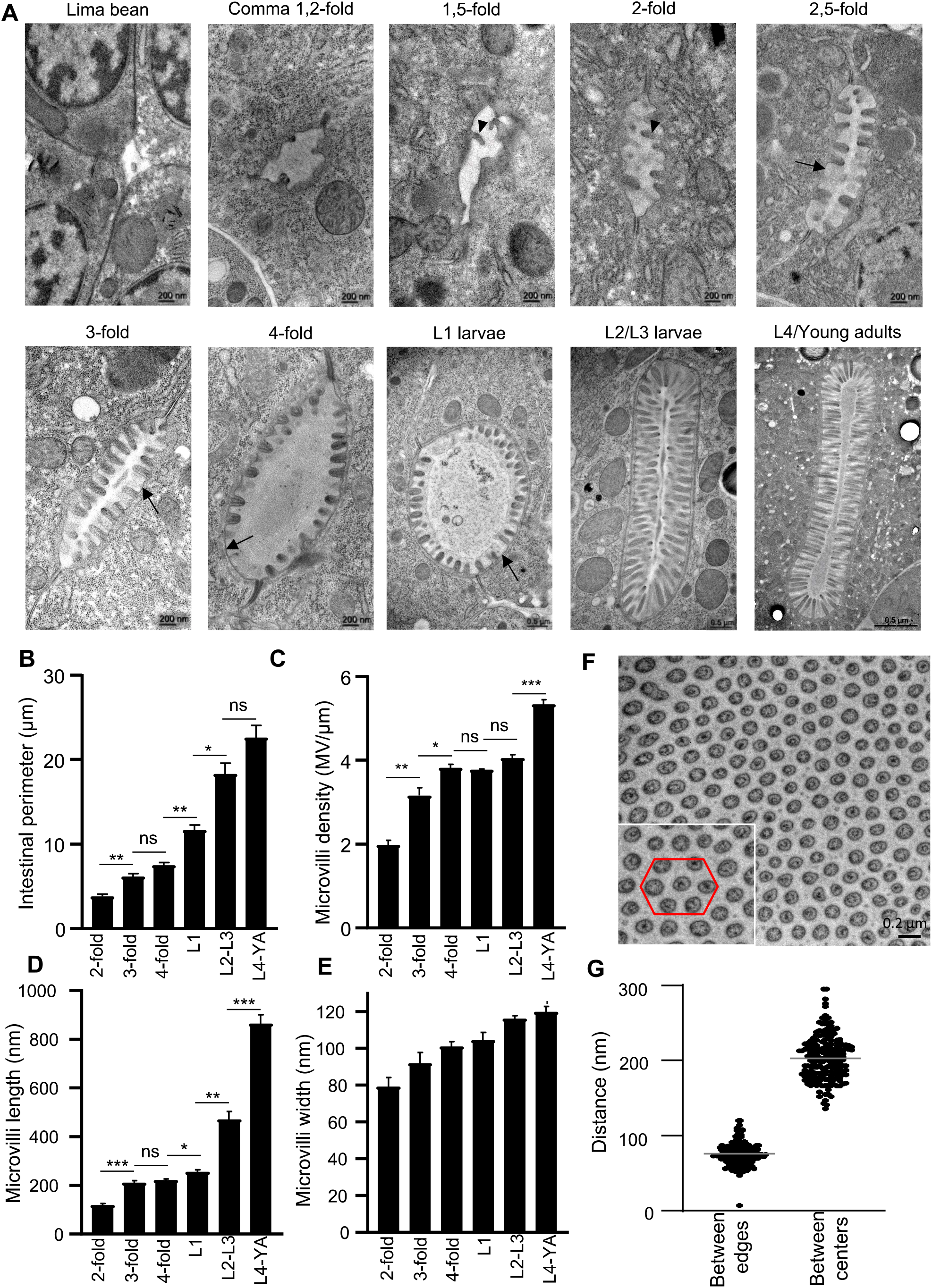
TEM analysis of the brush border. (A) Representative TEM images of the intestinal lumen at the *C.elegans* developmental stages indicated. (B-E) Quantification of the lumen perimeter (B) and microvilli density (C), length (D), and width (E) from TEM images. Histograms show the mean± SEM of the average of 3-13 slices (B), 3-13 lumen (C) and 6-29 microvilli (D-E) per sample from 3-5 worms at each developmental stage. Arrowheads, nascent microvilli; arrows, empty spaces between microvilli; YA, young adults. n.s., non-significant, * p<0,05, **p<0,01, ***p<0,001, unpaired t-test. (F-G) Transversal view of the brush border in adult worms. (G) The distance between microvilli edges and centres was calculated on 200 microvilli from (F).

### Dynamic recruitment of brush border components during C. elegans development

Expression profiling in mammalian enterocytes between the proliferative crypt and the terminally differentiated villus demonstrated a marked upregulation of actin cytoskeleton-related genes, including actin, ezrin, villin and espin (Chang et al., 2008, Mariadason et al., 2005). Notably, recent data in LLC-PKI cells showed a stepwise recruitment of EPS8 and IRTKS first, and then ezrin, during microvilli outgrowth (Gaeta et al., 2021). We hypothesized that a set of brush border components may specifically be recruited at the apical pole during brush border establishment *in vivo.* To test this, we performed a systematic analysis of the apical localization of endogenously tagged known brush border markers and putative new components, based on expression patterns as well as sequence or function homology with microvilli-associated mammalian proteins.

This led first to the identification of PLST-1, the ortholog of plastinl/fimbrin (Figs 2A and S2), which is one of the major F-actin organizing factor in mammalian cells brush borders, together with ezrin, villin and espin (Crawley et al., 2014a). While the ortholog of villin seems not to be localized at the brush border (Hunt-Newbury et al., 2007) and espin does not have a C. *elegans* ortholog, PLST-1 has been involved in cortical contractility in the C. *elegans* zygote (Ding et al., 2017) but has not been studied in the intestine yet. Second, the F-actin cross-linker FLN-2 (the ortholog of filamin A), that has been proposed to play a role in brush border maintenance in mammalian models (Zhou et al., 2014) was also localized at the enterocytes apical membrane in C. *elegans* (Figs 2B and S2).

**Figure 2.**
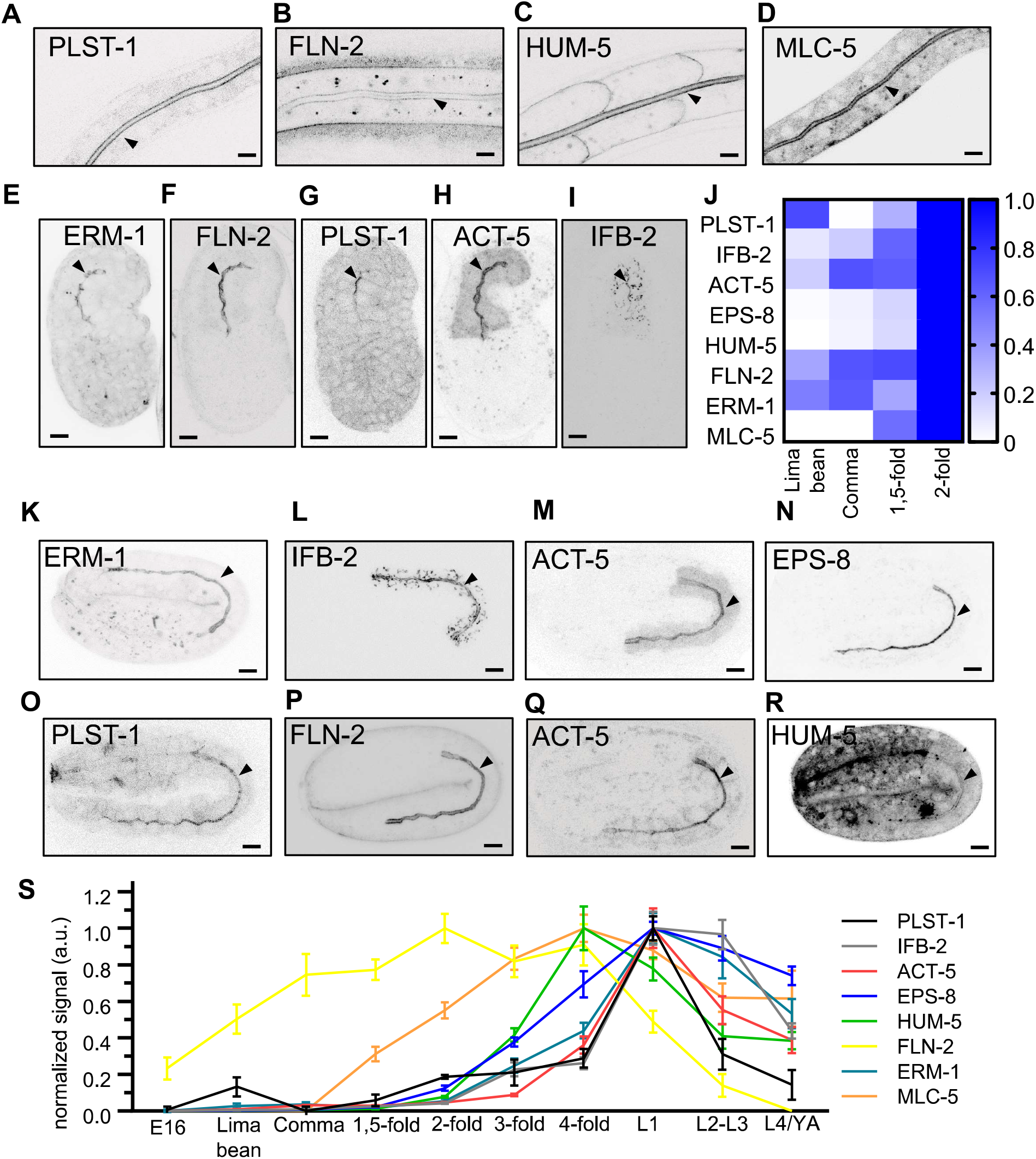
Brush border components are dynamically enriched at the apical membrane during microvilli assembly. (A-1) Representative images of the apical localization of GFP-tagged MLC-5, PLST-1 and ACT-5, mNG-tagged ERM-I, IFB-2 and HUM-5, and mVenus-tagged FLN-2 in Ll larvae (A-D) and lima bean embryos (E-1). (J) The absolute apical signal of the indicated markers was measured on at least 10 embryos at each developmental stage. The heatmap shows a focus on early brush border assembly steps where the maximum intensity was set at the 2-fold-stage. (K-R) Apical localization of the indicated markers at the 2-fold stage. (S) The absolute apical localization of the indicated markers was recorded as in (J) and normalized to the maximum expression for each marker. Data are mean± SEM. In all images, arrowheads show the apical plasma membrane of the intestinal cells. Scale bar, 5 μm.

In addition to these actin cytoskeleton organizing factors, many members of the myosin superfamily of actin motors have been localized to the brush border in mammalian cells (McConnell et al., 2011, Sauvanet et al., 2015). This superfamily comprises 12 classes of conventional and unconventional myosins, which function as multimers of heavy and light chains (Fili and Toseland, 2019). In the brush border, they have been shown to fulfil structural (e.g. MYO7b, MYH14), trafficking (e.g. MYO-la, −6) as well as contractile (non-muscle myosin NM2C) functions (Houdusse and Titus, 2021, Chinowsky et al., 2020). We found that a specific set of myosins accumulates at the enterocytes apex throughout C. *elegans* development: i) the unconventional heavy chain HUM-5 (the ortholog of human MYOld/g) which is also localized at the lateral membrane (Figs 2D, S2), but not the other members of this class, HUM-I and HUM-2 (Fig S3A-B); ii) the essential myosin light chain MLC-5 (Gally et al., 2009) (the ortholog of human MYLI/6) accumulated at the apical membrane of the enterocytes in both embryos and larvae (Figs 2D, S2), while MLC-4 was only weakly expressed in embryos (Fig S3C). Interestingly, we found that NMY-1 and NMY-2 (the orthologs of the non-muscle myosins NM2A/B and NM2B/C, respectively) (Fig. S3D-E), did not, or only very weakly for NMY-1, accumulate apically, which suggests that myosin-dependent contractility may be less crucial for microvilli assembly in C. *elegans* than in mammals (Chinowsky et al., 2020). These results suggest species-specific mechanisms or compensation between myosins, as shown before (Houdusse and Titus, 2021), and the need for systematic approaches to better characterize the conserved components of brush borders, for instance to identify putative intermicrovillar bridges molecules (e.g. protocadherins) (Crawley et al., 2014b) among the various cadherin-like proteins in the C. *elegans* genome (Loveless and Hardin, 2012).

To quantitatively assess the expression of these apically enriched factors during brush border establishment, we used photon counting detectors and quantified the absolute apical signal of endogenously tagged proteins at all C. *elegans* developmental stages (Figs 2S, S2 and S3F). Notably, we observed that a set of markers was already localized at the apical PM at the lima bean stage, before microvilli onset: ERM-1, FLN-2, PLST-1, ACT-5 (note that ACT-5 was exogenously expressed under its own promoter, because of the embryonic lethality of endogenously tagged strains), as well as the intermediate filament IFB-2, which appears slightly later (Figs 2E-J and S2). Then, we observed that the apical localization of these markers, as well as that of the actin polymerization/severing agent EPS-8, HUM-5 and MLC-5, dramatically increased concomitantly with microvilli assembly (from the 1.5-fold stage), most of them peaked between the 4-fold and LI stages and then decreased until adulthood (Fig. 2K-S and S2). The early accumulation of some of these factors at the apical membrane might reflect their importance for microvilli assembly, which is consistent with the requirement of ERM-I and ACT-5 (Gobel et al., 2004, MacQueen et al., 2005) and the direct relationship between G-actin apical availability and microvilli growth (Faust et al., 2019). As PLST-1 also accumulated before microvilli onset, it could also play a role in microvilli initial assembly *in vivo,* which is coherent with the disorganized terminal web and microvilli rootlets described in *Plsl* knockout mice (Grimm-Gunter et al., 2009), despite no obvious defect in *plst-1* C. *elegans mutants* at larval stages (not shown). Interestingly, we observed that FLN-2 displayed a shifted pattern, with an earlier peak that may suggest a specific role in microvilli establishment. Thus, an evolving set offactors might control microvilli building in C. *elegans:* a *pre-assembly* module, composed, at least, of ERM-1, ACT-5, PLST-1, FLN-2 and IFB-2 ; an *assembly module,* composed additionally ofEPS-8,HUM-5 and MLC-5 and, finally, a *mature module* that does not require FLN-2 (Fig. S5B).

### Super-resolution imaging of the brush border in vivo

According to the Rayleigh criterion 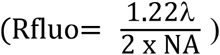, the optical axial resolution of the 405, 488 and 561 nm lasers is 176.5, 212.6 and 244.4 nm, respectively, using a NAl.4 objective, which is at the limit to resolve individual microvilli (~100 nm between edges,~200 nm between centers,~120 nm wide, Fig. 1E,G). To test this, we inserted three different tags at the C-terminal end of the brush border-specific factor ERM-1 by CRISPR-CAS9: Blue Fluorescent Protein (mTagBFP2/BFP, λ_Ex_ 381 λ_Em_ 445 nm) (Subach et al., 2011), mNeongreen (mNG,λ_Ex_ 506 λ_Em_ 517 nm) (Shaner et al.,2013) and wrmScarlet (wSc,λ_Ex_ 569 λ_Em_ 593 nm) (El Mouridi et al., 2017) and imaged them with a multi-detector and deconvolution-based super-resolution imaging system (see methods). We could easily visualize the regular alignment of microvilli with BFP and mNG tags, but it was less visible with the wSc fluorophore (Fig. 3A-B). In addition to individual microvilli, we could also precisely localize brush border markers along the microvilli long axis. Indeed, while ERM-I covered the whole microvilli length, the chloride intracellular channel 2 (CLIC-2) ortholog EXL-1 (Liang et al., 2017) accumulated at the tip ofmicrovilli (Fig. 3C) and the P-GlycoProtein related transporter PGP-1 at their base (Bidaud-Meynard et al., 2019). Of note, this method allowed to uncover localization differences between in-locus mNG-tagged and overexpressed GFP-tagged proteins (compare Figs 3C and S4A), as already reported for E-cadherin in the same tissue (Cordova-Burgos et al.,2021). Individual microvilli were similarly visualized using Random Illumination Microscopy (Mangeat et al., 2021), but not using conventional confocal or Stimulated-emission-depletion (STED) microscopes, probably because of the depth of the intestine inside the nematode body (~15μm) (Fig S4B). The brush border could also be imaged transversally (compare Figs 3D and 1F). Hence, the combination of a specific super-resolution imaging system, appropriate fluorophores and endogenous expression allows the precise visualization of microvilli *in vivo* in C. *elegans* intestine.

**Figure 3.**
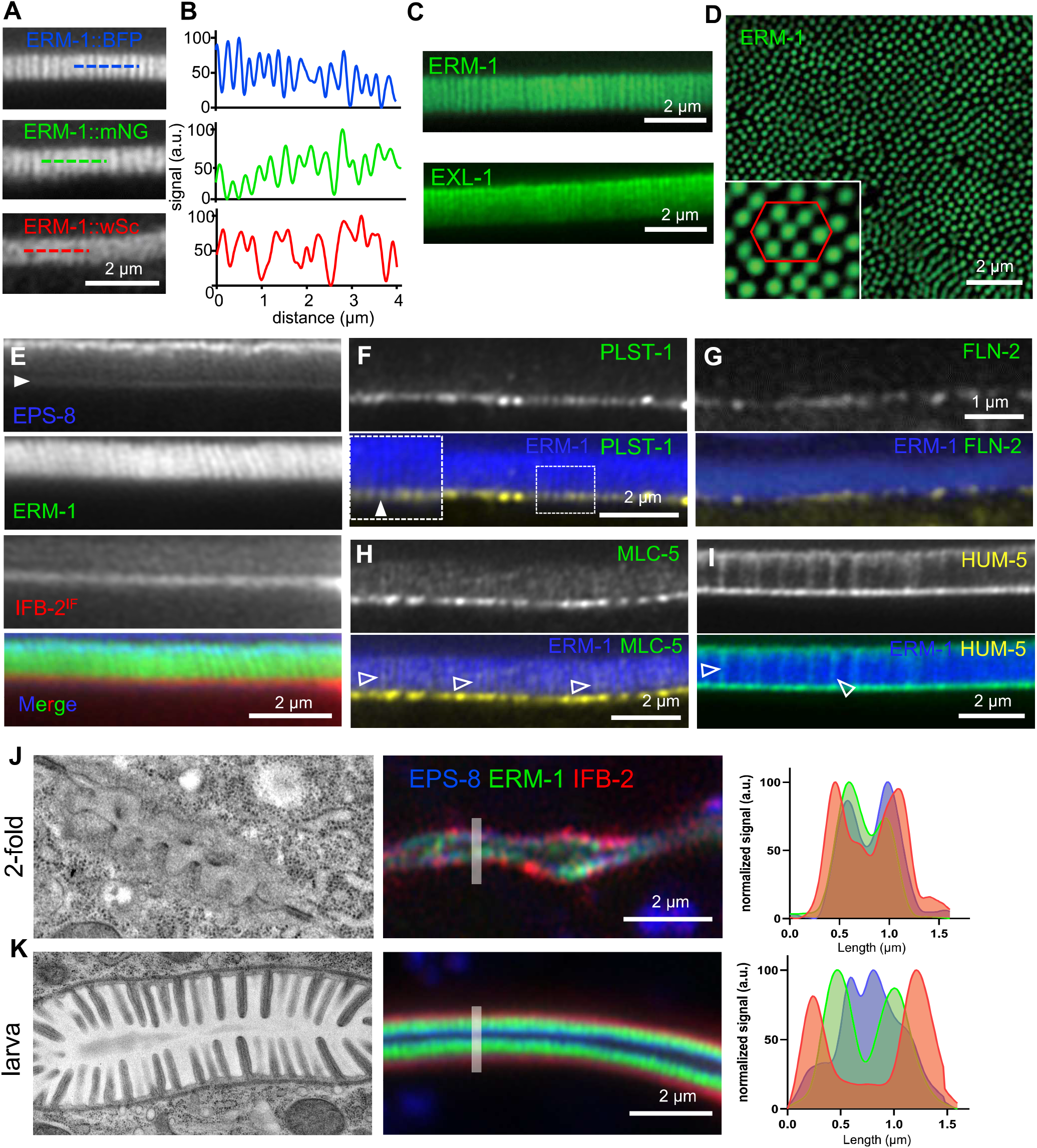
Super-resolution imaging of the brush border. (A-B) Super-resolution images ofERM-I endogenously tagged with BFP, mNG and wSc in C. *elegans* young adults. (B) represents the normalized intensity profile along a 4 μm-long dashed line, as represented in (A). (C) Super-resolution images ofERM-1::mNG or EXL-1::mNG in young adults. (D) Transversal super-resolution image of the brush border performed on a young adult C. *elegans* strain endogenously expressing ERM-1::mNG. The red hexagon indicates the putative hexagonal packing ofmicrovilli. (E-I) Representative super-resolution images of the indicated microvilli markers endogenously tagged with mNG (ERM-1, HUM-5,), GFP (PLST-1, MLC-5), BFP (EPS-8), mVenus (FLN-2) or wSc (IFB-2) in young adults. Insert, higher magnification of the ROI depicted by the dotted line. Closed and open arrowheads show the colocalization between ERM-1 and the indicated markers at the base and along the microvilli, respectively. (J-K) Left TEM images show the shape of the brush border at the corresponding developmental stage. Right: super-resolution images of the brush border in 2-fold embryo and L2 larvae co-expressing EPS-8::BFP, ERM-1::mNG and IFB-2::wSc. Right histograms correspond to the signal intensity profile of the three markers along the line depicted on the pictures.

We then used these tools to study the (co)localization of known and newly identified apical markers in adult worms, as we have done before for ERM-1 and ACT-5 (Bidaud-Meynard et al., 2019). Using a strain co-expressing endogenously tagged versions of the three classical microvilli markers ERM-1, EPS-8, and IFB-2, we observed that ERM-1 localized along the whole microvilli but not in the terminal web (Fig. 3E). EPS-8 accumulated at the tip of the microvilli, where it partially colocalized with ERM-I, and was also found marginally at the terminal web vicinity, as observed before by immuno-EM (Croce et al., 2004) (Fig. 3E). Finally, we could resolve in some worms the tiny difference between ACT-5, which localized along and at the basis of the microvilli, and the endotube marker IFB-2 (Geisler et al., 2019, Bossinger et al., 2004), with which it composes the terminal web (Fig. S4C).

Notably, we found that PLST-1 localized at the bottom of the microvilli (Fig. 3F), with a doted pattern different from the linear terminal web pattern (IFB-2 in Fig 3E). This localization is consistent with that ofPlastin-1 in mouse jejunum sections and its proposed role in anchoring microvillar actin rootlets to the terminal web (Grimm-Gunter et al., 2009). While FLN-2 was hardly detectable in adult worms, we observed in LI larvae that FLN-2 localized at the basis of microvilli (Fig. 3G), alike MLC-5, which also partly decorated the whole microvilli (Fig. 3H). Finally, we found that HUM-5 accumulates at both the basis and the tip of, and localizes slightly along microvilli (Fig. 3I), a similar pattern to that described in mouse intestine (Benesh et al., 2010). Thus, our novel approach allowed to visualize *in vivo* the expression and the precise localization of several proteins of the brush border including structural proteins and molecular motors.

Since factors needed to build the microvilli are concomitantly recruited to the apical pole (Fig. 2), we finally asked whether super-resolution imaging could resolve the change in their relative microvillar position during brush border assembly. Line scans showed that ERM-1, EPS-8 and IFB-2 colocalized at the beginning of microvilli assembly (2-fold stage) and progressively moved away to end up with IFB-2 and EPS-8 contralaterally positioned and surrounding ERM-1 (Figs 3J-K and S4D).

### Analysis of brush border markers dynamics during microvilli assembly

To test whether the progressive accumulation of brush border components at the apical membrane implies a dynamic behaviour during microvilli building (Fig. 2S), we analysed the dynamics of ERM-I during and after the establishment of the brush border *using.fluorescence recovery after photobleaching* (FRAP) experiments. While ERM-I was very dynamic during microvilli assembly (1.5-fold embryo) it became surprisingly extremely stable in established brush border (adult worm), with little recovery even after>15 minutes (Figs 4A and S5A), confirming recent observations (Ramalho et al., 2020, Remmelzwaal et al., 2021). Systematic analysis of ERM-I fluorescence recovery throughout C. *elegans* development confirmed that ERM-1 dynamics progressively decreased concomitantly with brush border assembly and became almost static in larvae and adults (Fig. 4B, F-G). To confirm this, the dynamics of other structural components of the brush border was analysed during microvilli initial assembly (Comma/1.5-fold), maturation (LI larvae) and in adult worms; note that due to embryo fast movements from the 2-fold stage to the end of embryogenesis, these developmental stages could not be investigated. Like ERM-I, EPS-8 was also very dynamic during microvilli assembly but became highly stable in maturating and mature microvilli (Fig. 4C, F-G). ACT-5 also displayed a dynamic, albeit of a lower extend, behaviour, that persisted until LI larvae (Fig. 3D, F-G), in agreement with F-actin mobile fractions in Caco-2 cells (~60%) (Waharte et al., 2005), to finally become stable at adulthood. Conversely, the intermediate filament IFB-2 displayed a more stable behaviour at every developmental stage, which reflects its anchoring role for growing microvilli (Grimm-Gunter et al., 2009, Geisler et al., 2019).

**Figure 4.**
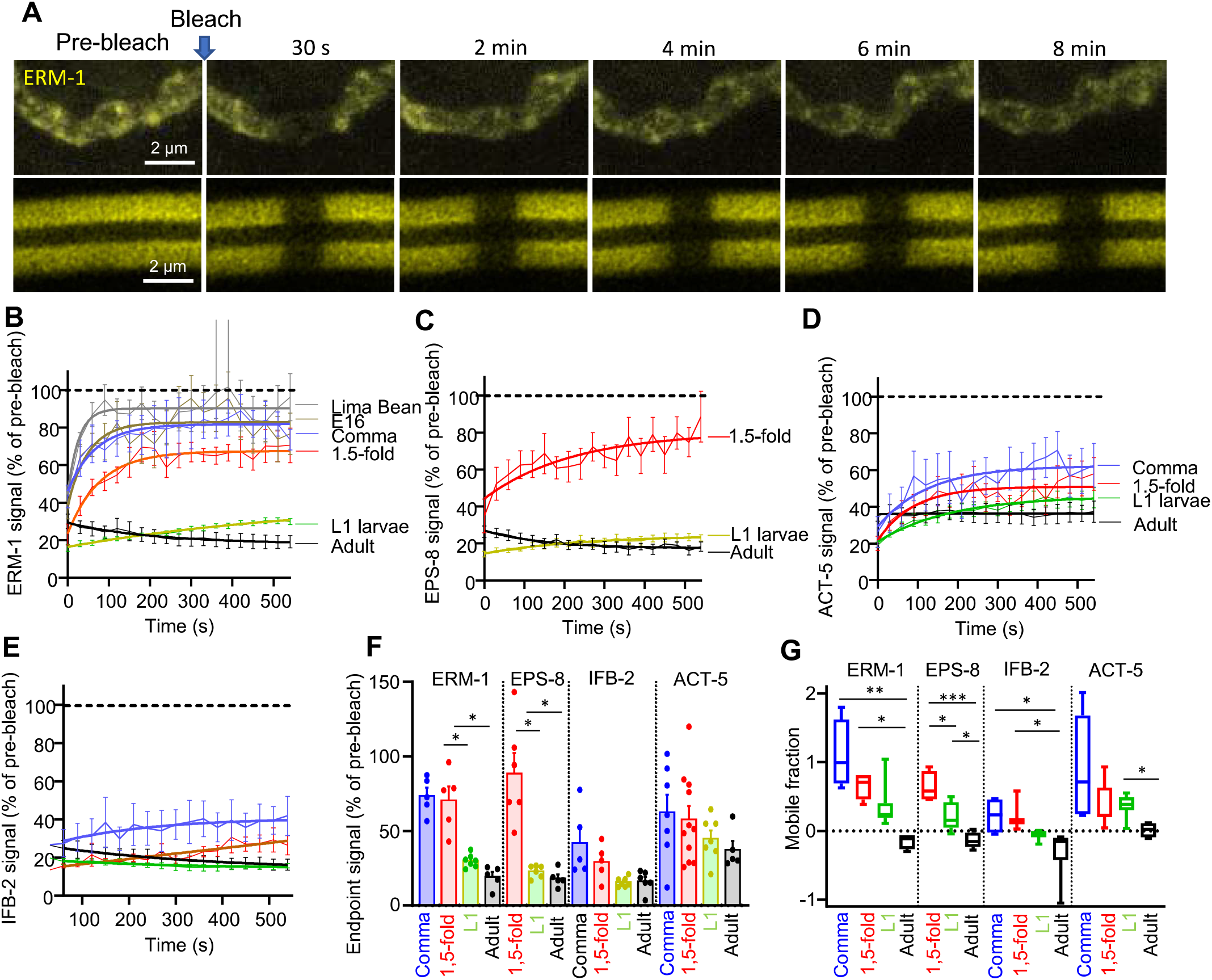
Brush border components dynamics during microvilli assembly. (A-B) ERM-1::mNG was bleached in a 1.5-fold embryo and an adult worm, and fluorescence recovery was observed every 30 s. (B-E) Quantification of the signal recovery after bleaching of ERM-I ::mNG (B), EPS-8::mNG (C), ACT-5::GFP (D) andIFB-2::mNG (E), measured every 30 s on 5-11 worms at the indicated developmental stages. Thin lines represent the mean± SEM of signal recovery, bold lines represent one-phase association non-linear regression fitting curves. (F-G) Analysis of the FRAP experiments performed in (D-E) at the comma, 1.5-fold, LI larva and adult stages. (F) Shows the % of the pre-bleached signal (mean of two timepoints) recovered at the endpoint (540 sec) and (G) shows the mobile fraction. Histograms show the mean± SEM, dots in (F) represent individual worms. The difference between variance was calculated using ANOVA, *p<0,05, **p<0,01, ***p<0.001.

Thus, these results enlightened that mature microvilli adopt a stable steady state *in vivo,* which is consistent with the notion that microvilli might be considered more as stereocilia than evanescent F-actin-based structures like filopodia. The maturation status of the brush border might be a key consideration that would help to reconcile conflicting data of the literature. Indeed, probably immature microvilli in non-polarized cells seem to be more dynamic, i.e. life-cycle of~12 min in A6 cells (Gorelik et al., 2003), with an intense actin treadmilling (half-time recovery of ezrin of~30 s) (Garbett and Bretscher, 2012) but which were found to last up to 12 h in mature brush borders (Meenderink et al., 2019). This high stability could contribute to explain their uniform length and highly ordered organisation in the human intestine (Crawley et al., 2014a).

In conclusion, this new multi-imaging approach allowed to image the precise localization of brush border markers at the microvilli level *in vivo* and to study the dynamic recruitment of microvilli components during the development of the brush border. This new methodology will be instrumental to address the many questions remaining to understand microvilli assembly and maturation, notably on the full set of factors required for microvilli growth and maintenance, the principles that govern microvilli size, packing and organization or the motility of microvilli *in vivo.* It will be also essential to understand the pathophysiology ofdiseases affecting the brush border, such as Microvillus inclusions disease (Bidaud-Meynard et al., 2019), Crohn’s (VanDussen et al., 2018) and celiac (Tye-Din and Anderson, 2008) diseases, or pathogen infections (Scott et al., 2004, Lauwaet et al., 2004), as well as intestinal aging.

## Materials and methods

### *C. elegans* strains and maintenance

Strains were maintained under typical conditions as described (Brenner, 1974). CRISPR-CAS9-genome edited mTagBFP2, mNeonGreen and mScarlet-tagged proteins were generated at the « Biologie de *Crenorhabditis elegans* » facility (Universite Lyon 1, UMS3421, Lyon, France). The strains used in this study are listed in Table S1.

### *in vivo* confocal imaging in *C. elegans*

For *in vivo* imaging, C. *elegans* larvae were mounted on a 10% agarose pad in a solution of 100 nm polystyrene microbeads (Polysciences Inc.) to stop worm movement. Embryos were mounted on a 2% agarose pad with a mix of bacteria and M9 medium (localization) or M9 only (live imaging). Single confocal slices of the anterior intestinal cells or stacks of the whole intestine were performed on adults/larvae and whole embryos, respectively, using a Leica SP8 (Wetzlar, Germany) equipped with a 63X, 1.4 NA objective (LAS AF software) or a super-resolution Zeiss LSM880-Airyscan (Oberkochen, Germany) equipped with a 63X, 1.4 NA objective (Zen Black software). Quantitative recording of the apical localization of brush border markers was performed on the Leica SP8 microscope using the photon counting function of HyD hybrid detectors and image accumulation (Fig. S3). For embryos, stacks were reconstructed using the max intensity Z-projection function of Fiji software (https://imagej.net/Fiji). All images were examined using Fiji software.

### TEM

Samples were subjected to high-pressure freezing followed by freeze substitution, flat embedding, targeting, and sectioning using the positional correlation and tight trimming approach, as described previously (Bidaud-Meynard et al., 2019). Each embryo or larva was sectioned in 5-10 different places, every 5-7 μm, to ensure that different intestinal cells were observed. Ultrathin sections (60-70 nm) were collected on formvar-coated slot grids (FCF2010-CU, EMS) and observed using a JEM-1400 transmission electron microscope (JEOL, Tokyo, Japan) operated at 120 kV, equipped with a Gatan Orius SC 1000 camera (Gatan, Pleasanton, USA) and piloted by the Digital Micrograph program.

### Fluorescence recovery after photobleaching (FRAP)

FRAP experiments were performed using the Zeiss LSM880-Airyscan on a rectangle ROI of 1,3 μm width crossing the apical PM with 100% 488 nm laser power, 10-20 iterations and recovery was measured every 30 s for 10 to 15 min. Post-FRAP images were analysed using the Fiji software. The mean fluorescence intensity of the bleached ROI was normalized for photobleaching by recording the intensity of the same ROI on a non-bleached region and cytoplasmic background was subtracted on each frame. Finally, the % recovery was calculated on each timeframe by comparing the normalized signal intensities with the mean of two timepoints before bleach. Curve fitting was performed with one-phase association non-linear regression analysis using Graphpad Prism 9 software. The mobile fraction (Mt) was calculated with the following equation: 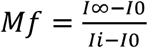 where I_∞_ is the signal intensity at the endpoint plateau phase, I_0_ is the mean of two pre-bleached signal intensities and Io is the signal intensity at the first post-bleach timepoint.

### Quantification

Micrographs were analysed using Fiji software and were representative of all the sections observed. Microvilli (length, width, density) and lumen perimeter were quantified on at least 6-13 TEM images per sample (n2:3, by developmental stage).

For the quantitative measurement of the apical localization of brush border markers, a maximum intensity projection was performed using Fiji, and the signal density was quantified by measuring the mean fluorescence signal along a segmented line covering the whole intestine (El6 to 2-fold embryos) or visible part of the anterior intestine (3-fold to adults). The signal measured was then corrected for fluorescence accumulation and normalized.

### Statistical analysis

Results are presented as mean ± SEM of the number of independent experiments indicated in the legends, and scattered dots represent individual worms. p-values were calculated by two-tailed unpaired student’s t-test or one-way ANOVA, as indicated in the figure captions, and a 95% confidence level was considered significant. Normal distribution of data and homogeneity of variances were validated using the Shapiro-Wilk and the Bartlett (for ANOVA) or F (for student) tests, respectively. Mann-Whitney or Kruskal-Wallis tests were used for calculating the P-values between two or multiple non-normal distributions, respectively, and Dunnett’s tests was used for multiple distributions with non-homogenous vanances.

## Acknowledgements

We thank Marc Tramier and Stephanie Dutertre for their advice on fluorescence quantification and super-resolution imaging, respectively, as well as Matis Soleilhac for the initial analysis of expression patterns. We also thank Michel Labouesse, Junho Lee and Ronen Zaidel-Bar for strains as well as Celine Burckle and Guillaume Halet for helpful discussions. Some strains were provided by the CGC, which is funded by NIH Office ofresearch Infrastructure Programs (P40 OD010440; University ofMinnesota, USA). We are grateful to Maite Carre-Pierrat who performed CRISPR-CAS9 endogenous tagging at the Biology of *Caenorhabditis elegans* Facility, Universite Lyon 1, UMS3421, France. Photonic and electron microscopy imaging were performed at the Microscopy Rennes imaging Center (MRiC, Biosit, Rennes, France), member of the national infrastructure France-Bioimaging supported by the French National Research Agency (ANR-10-INBS-04).

## Funding

This work was supported by the European Union’s Horizon 2020 research and innovation program under the Marie Sklodowska-Curie grant agreement 844070 to ABM, Defis scientifiques de l’Universite Rennes 1 (17CQ436-S0) to ABM and GM, Ligue Regionale Contre le Cancer (22, 29, 35, 41, 72, 85) and the Fondation maladies rares (169608 and EXM-2019-1013) to GM. GM laboratory also received institutional funding from the CNRS and the Universite de Rennes 1.

## Authors contribution

Conceptualization: A.B.M., G.M.; Methodology, Validation, Formal analysis, Investigation, Data curation: A.B.M., F.D., O.N., A.P., G.M.; Ressources: S.K.S., C.P., F.R.; Writing - original draft: A.B.M. G.M.; Writing - review & editing: A.B.M., O.N., A.P., G.M.; Supervision: G.M; Funding acquisition: A.B.M., G.M.

## Conflict of interest

The authors declare no conflict of interest.

---

**Figure S1.**
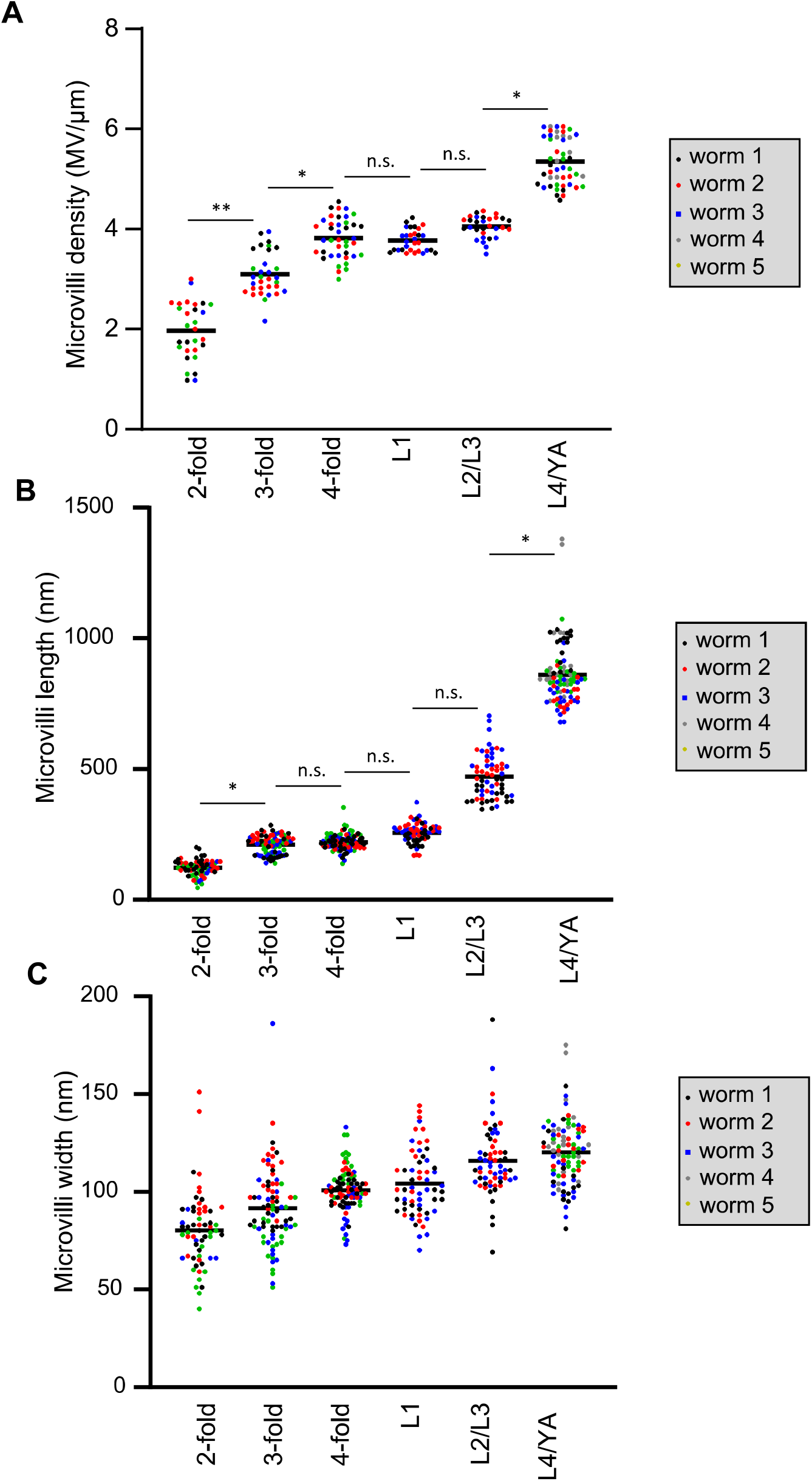
Individual values of brush border measurements by TEM, related to Fig. 1. Colorized dots represent individual worms at the indicated developmental stages. Bar is the grand mean of all the measurements. Microvilli density was measured on 3-13 slices/worm, microvilli length and width on 6-29 microvilli/worm. N.s., non-significant, *p<0,05, **p<0,01, Mann-Whitney test (A-B) and unpaired t­ test (C).

**Figure S2.**
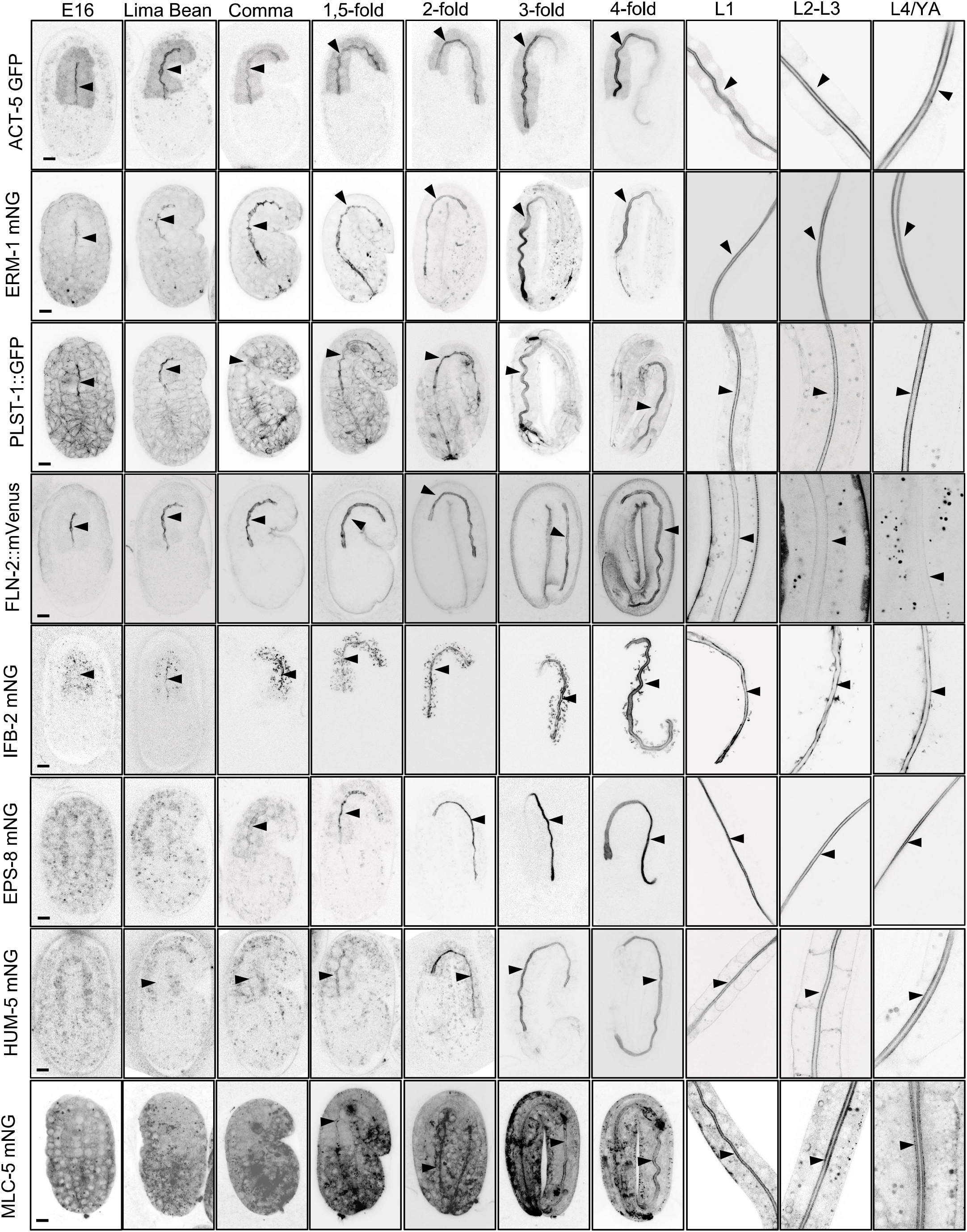
Systematic analysis of brush border markers during *C. elegans* development. Representative confocal images of the endogenously tagged markers indicated (except ACT-5::GFP). Arrowheads show the intestinal cells apical PM. Scale bar is 5 μm.

**Figure S3.**
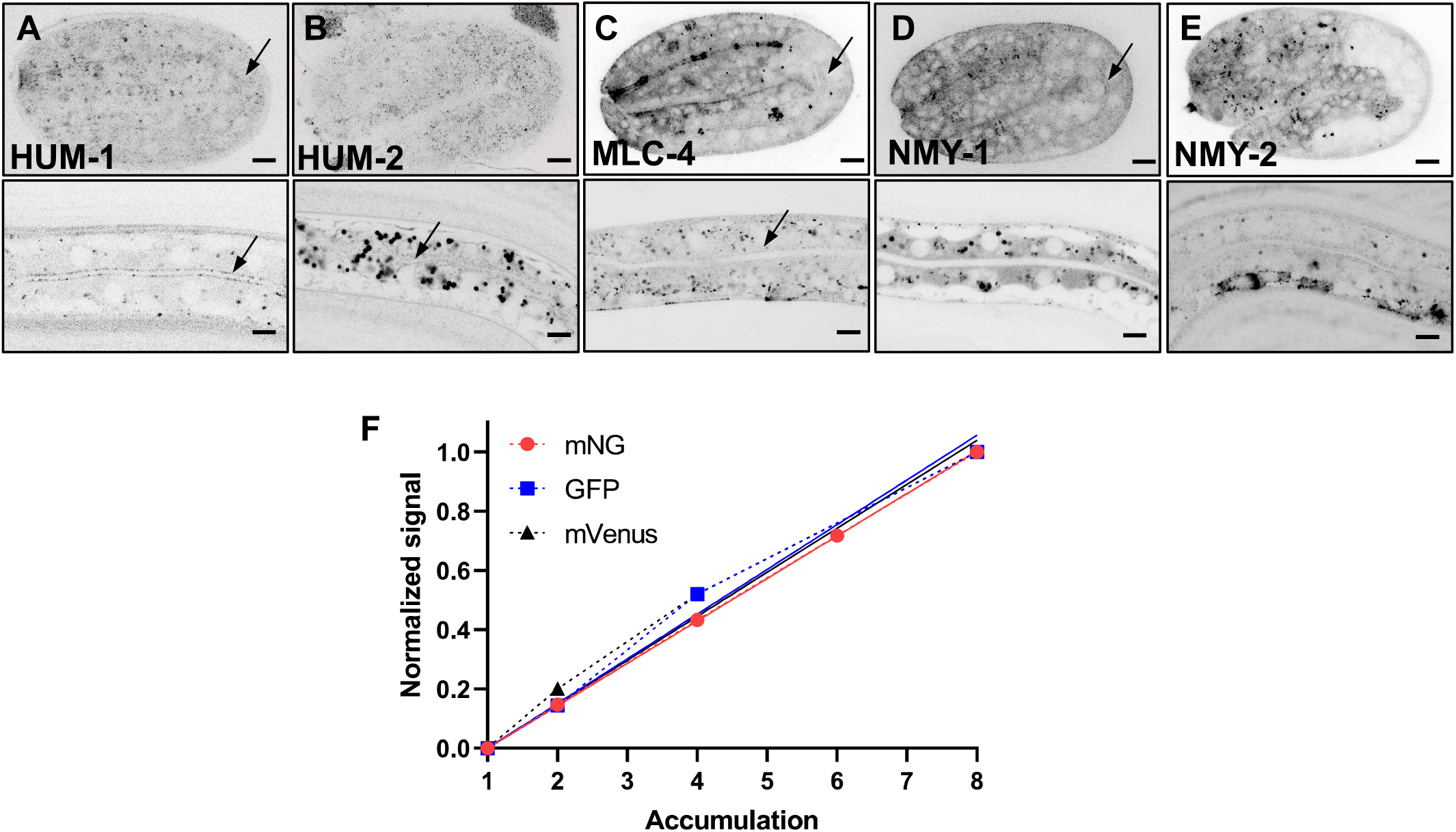
Systematic analysis of brush border markers during C. elegans development. (A-E) Representative confocal images of C. *elegans* strains expressing endogenously tagged versions of the indicated markers, which showed no apical accumulation during C. *elegans* intestine development. (F) Control of the quantitative assessment of brush border markers arrival at the apical PM. Accumulation of ERM-1::mNG, PLST-1::GFP and FLN-2::mVenus signal linearly increases with image accumulation. Scale bar, 5 μm.

**Figure S4.**
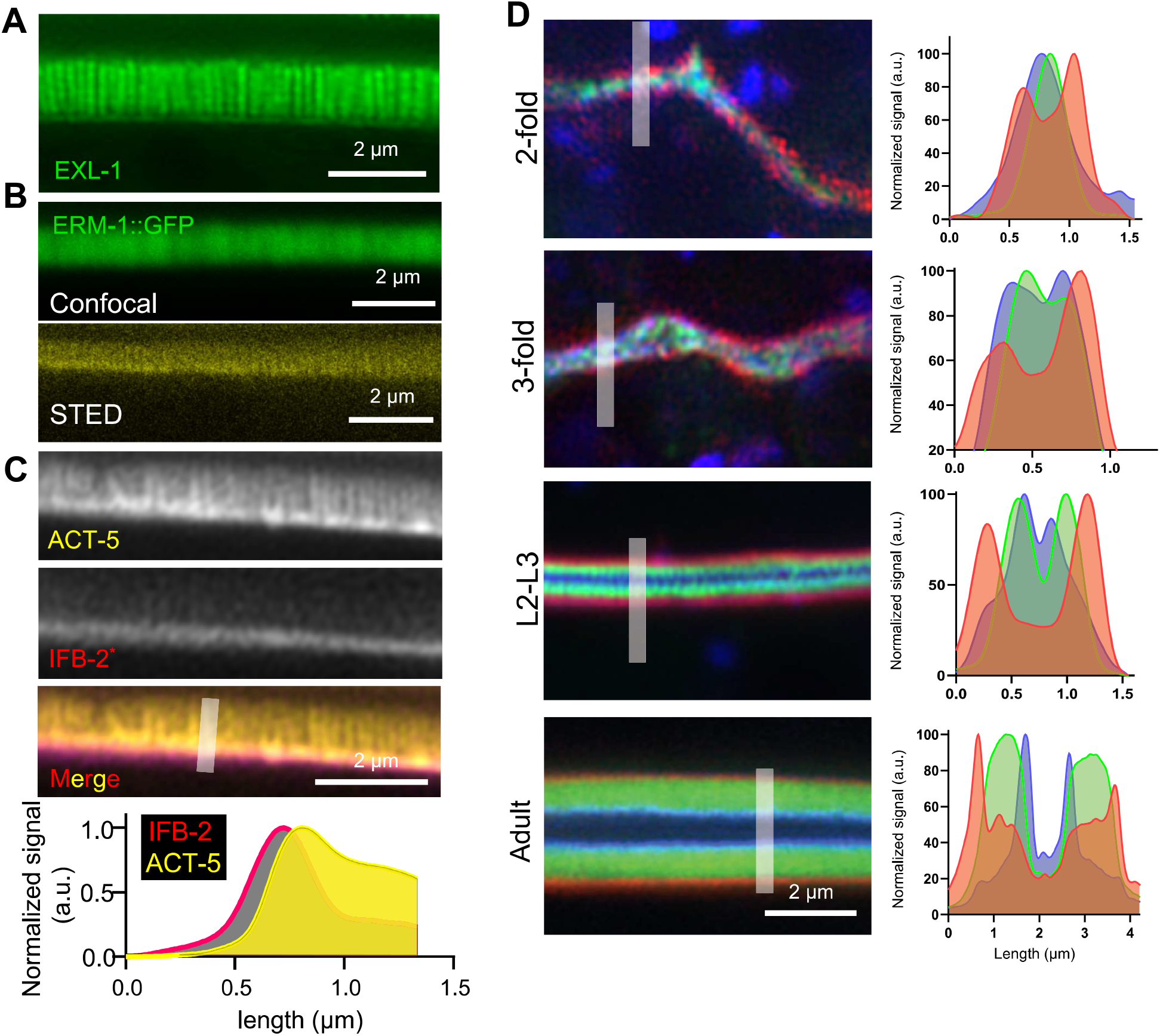
Super-resolution imaging of brush border markers *in vivo.* (A) Representative super­ resolution image of exogenously expressed EXL-1::GFP. (B) ERM-1::GFP was imaged in adult worms using the indicated microscopes. (C) Super-resolution images of a C. *elegans* adults co-expressing ACT-5::GFP and IFB-2::wSc. Bottom panel shows a normalized intensity profile along the line depicted in grey. (D) Representative images of the localization of endogenously tagged EPS-8::BFP, ERM-1::mNG and IFB-2::wSc in C. *elegans* at the indicated developmental stages. Right panels show an intensity profile of the three markers along the line depicted in left panels. Scale bars, 2 μm.

**Figure S5.**
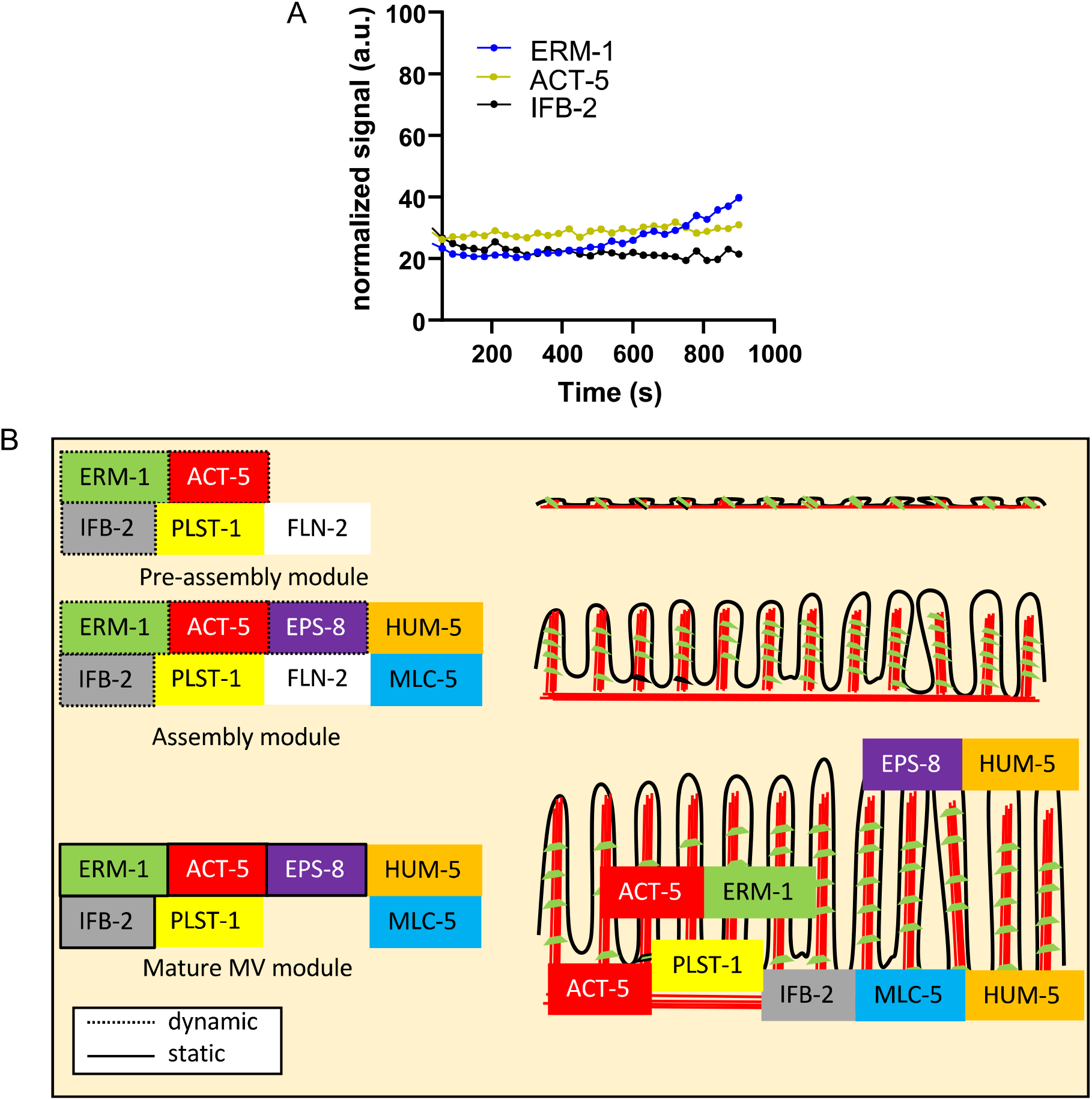
Dynamic recruitment of brush border components. (A) Longer measurement of brush border components dynamics in adult worms. The curves show the recovery ofERM-1::mNG, IFB-2::mNG and ACT-5::GFP signal every 30 s after photobleaching, measured for an extended time, n= l for each marker. (B) Model of brush border assembly *in vivo* in C. *elegans.* Microvilli are built from a preformed *pre-assembly module* and grow through the dynamic recruitment of brush border components, which become highly stable in the mature brush border.

**Table S1.**
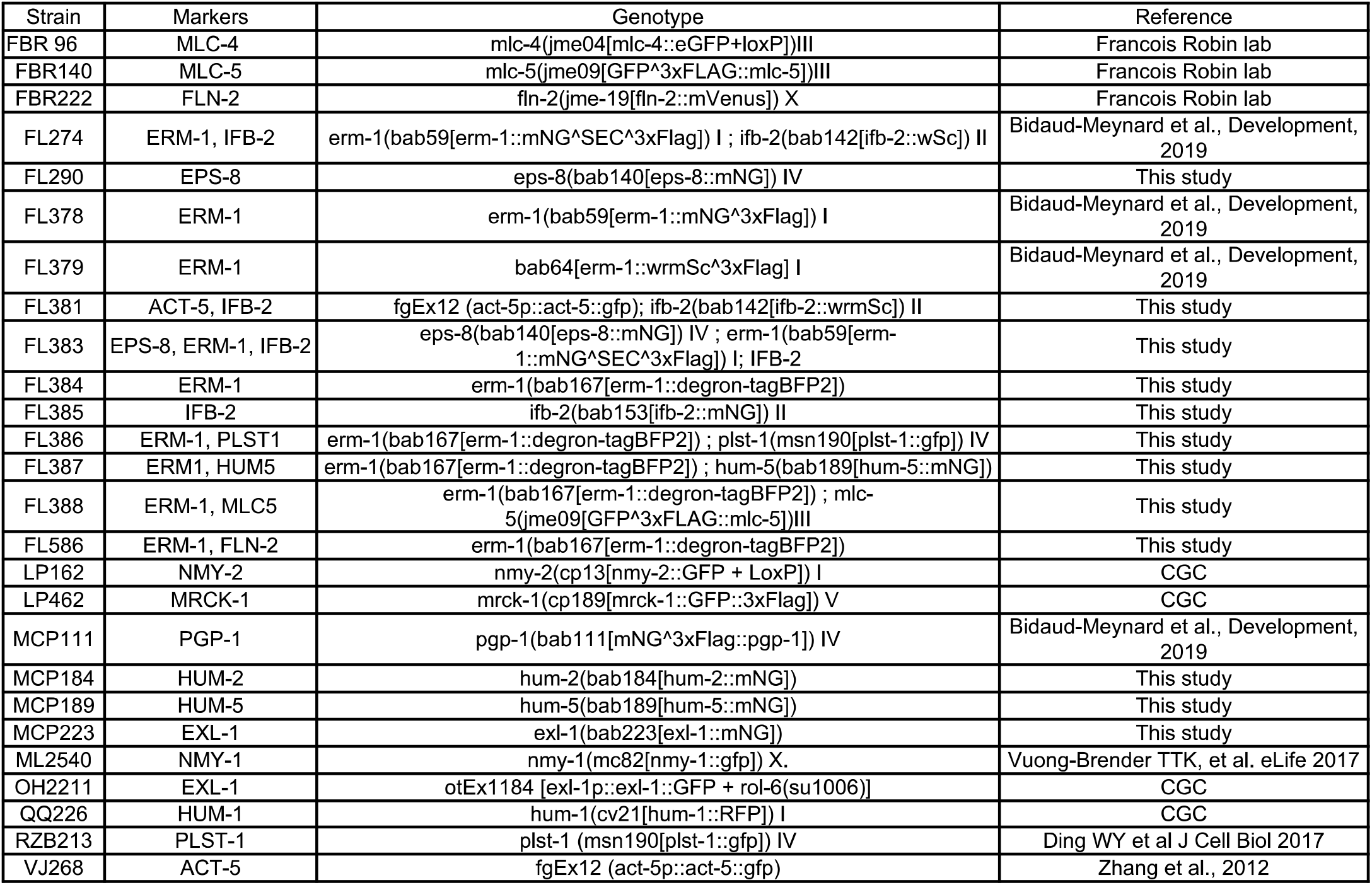
*C. elegans* strains used in this study.

## Notes

### Competing Interest Statement

The authors have declared no competing interest.

